# Damaging Mutations are Associated with Diminished Motor Skills and IQ in Children on the Autism Spectrum

**DOI:** 10.1101/141200

**Authors:** Andreas Buja, Natalia Volfovsky, Abba Krieger, Catherine Lord, Alex E. Lash, Michael Wigler, Ivan Iossifov

## Abstract

In individuals with Autism Spectrum Disorder (ASD), de novo mutations have previously been shown to be significantly correlated with lower IQ, but not with the core characteristics of ASD: deficits in social communication and interaction, and restricted interests and repetitive patterns of behavior. We extend these findings by demonstrating in the Simons Simplex Collection that damaging de novo mutations in ASD individuals are also significantly and convincingly correlated with measures of impaired motor skills. This correlation is not explained by a correlation between IQ and motor skills. We find that IQ and motor skills are distinctly associated with damaging mutations and, in particular, that motor skills are a more sensitive indicator of mutational severity, as judged by the type and its gene target. We use this finding to propose a combined classification of phenotypic severity: mild (little impairment of both), moderate (impairment mainly to motor skills) and severe (impairment of both).

## Introduction

Autism Spectrum Disorder (ASD) is a neuropsychiatric disorder, conventionally characterized by core phenotypes, including persistent deficits in social communication and interaction, and restricted, repetitive patterns of behavior (American Psychiatric Association, 2013). Genetics is a strong determining factor, as evidenced by the high rate of concordance between identical twins, elevated sibling risk, consistent enrichment of autism-associated genes in certain biological pathways and neurodevelopmental periods, and the presence of genetic ‘signatures’. These signatures include a significantly increased incidence of likely gene-damaging de novo small and large scale mutations (Iossifov et al. 2014, Sebat et al. 2007, Sanders et al. 2015, De Rubeis et al. 2014, Levy et al. 2011), and the preferential transmission of such variants to the affected child (Levy et al. 2011, Iossifov et al. 2015, Krumm et al. 2015)

Although the core phenotypes form the consensus clinical signature of ASD, such children also have a wide range of other phenotypes and comorbidities (American Psychiatric Association, 2013). A wealth of phenotypic data on autistic individuals is found in the Simons Simplex Collection (SSC), an archive of samples from ‘simplex’ families, where only one child is affected, and may include one or more unaffected children (Fischbach & Lord, 2010). The SSC samples have been the source for the discovery of many candidate causal de novo variants. The richly documented and quantified phenotypic variables in the SSC provide an excellent opportunity to correlate variants with phenotypes. In an earlier effort, we and others have shown that ASD individuals with low nonverbal IQ (nvIQ) have a significantly increased incidence of damaging de novo mutation (Iossifov et al. 2014, O’Roak et al. 2012). These observations support the hypothesis that damaging de novo mutations may have broader neurological effects than ASD alone.

Wishing to test this hypothesis further, we looked for further correlations between phenotypes and damaging de novo (dn) mutations. While IQ reflects some aspects of cognitive ability, there are other fundamental manifestations of altered neurological function. Indeed, neurodevelopmental delay, such as age of first walking, is often the first presenting symptom in autism (Landa and Garrett-Meyer 2006, Provost et al. 2007). The delay in this milestone may be more generally a reflection of diminished motor skills (MS), a well-documented feature of ASD (e.g., Paquet et al. 2016). Some have argued that motor impairment should be included among core ASD features (e.g., Mosconi and Sweeney 2015, Hilton et al. 2012, Fournier et al. 2010, Mostofsky et al. 2007, Dziuk et al. 2007). Thus we decided to look for correlations between MS and dn mutations, in particular those that are ‘likely gene disrupting’ (or LGDs: nonsense, frame shift and splice site altering).

Motor skills of most affected children in the SSC have been evaluated by the Developmental Coordination Disorder Questionnaire (DCDQ) and, for young children, also by the Vineland Adaptive Behavior Scales (VABS-II). Using these scores, we find dn LGD mutations correlate with MS at least as strongly and significantly as with nvIQ. Statistical significance of this correlation is found not only for the total DCDQ and VABS-II scores, but also for their subcomponents, including fine and gross motor skills, as well as for related variables such as delay in the developmental milestone “age of first walking”. Moreover, significance of the correlations increases when we weight a dn LGD mutation by evidence that its target gene is under strong purifying selection in humans, or that it is a member of certain functional classes. We extend our observations even further by including an analysis of missense mutations, in which correlation to MS (much less so IQ) becomes evident when these mutations are additionally weighted by predictions of deleterious effect.

While IQ and MS are correlated with each other, they each correlate with damaging dn mutations even after either is adjusted for the other. Although MS and IQ significantly correlate with the severity of the core ASD phenotypes, we observe no consistently significant correlation between damaging dn events and core ASD. This finding suggests that the source of social cognitive impairment in ASD may be variants in genes not experiencing particularly strong negative selective pressure, and therefore not entirely rare in the human population.

## Results

### Motor skills in those with de novo LGDs

Motor skills are scored in the DCDQ as a fifteen item parent questionnaire (on a 5-point Likert scale where higher values indicate higher achievement) that assesses the child’s ability for fine and gross motor skills (Schoemaker et al. 2006). (See the Appendix for more detailed descriptors.) The total DCDQ score and three summary subscores are available for 87% of the exome-sequenced affected children in the SSC. The subscores of the DCDQ are “control during movement”, “fine motor/handwriting”, and “general coordination.” Though not standardized for age, it is negligibly age dependent (S3 in the supplement). Additional measures of motor skill development are found in various instruments used for evaluating the affected children, in particular the Vineland-II Motor Skill Domain for young children, with a main scale and two subscales: fine and gross motor skills. In addition, a measure of coordination difficulty is found in the Social Responsiveness Scale (SRS): item 14, which asks for problems with being ‘well-coordinated’ (on a severity scale 0–3). Finally, the Autism Diagnostic Instrument (ADI-R) has a milestone variable “walked unaided, age” (item 5), and a variable “articulation at age 5” (item 32) that provides one measure of development of motor control of speech.

We examined correlations of the phenotypic features with the number of dn LGDs per child (typically 0 or 1, some 2, few 3) and displayed the strengths of their one-sided p-values graphically as seen in Figure 1, column A (see Materials and Methods for details). The DCDQ and VABS measures, as well as nvIQ, are skill levels, hence expected to be negatively correlated with dn LGD mutations (shown red). Three measures (shown in the bottom rows of the figure) are inverse skill measures, “age at first walking”, “speech articulation at age 5” and (problems with) “well coordination”, hence expected to be positively correlated with dn LGD mutations (shown blue). The correlations of all these measures with our primary measure of genetic damage are in the expected directions. The absolute correlations between genotype and MS measures tend to be low, on the order of 0.1 (Figure 1’ in supplement). However, due to the large size of available SSC data (n=2,120), these correlations are statistically significant (p=1.5E-4 for “total DCDQ”). Affected children with dn LGDs have decreased gross and fine motor skills, as well as delayed motor development, compared to affected children without these observed dn LGDs.

**Figure 1.**
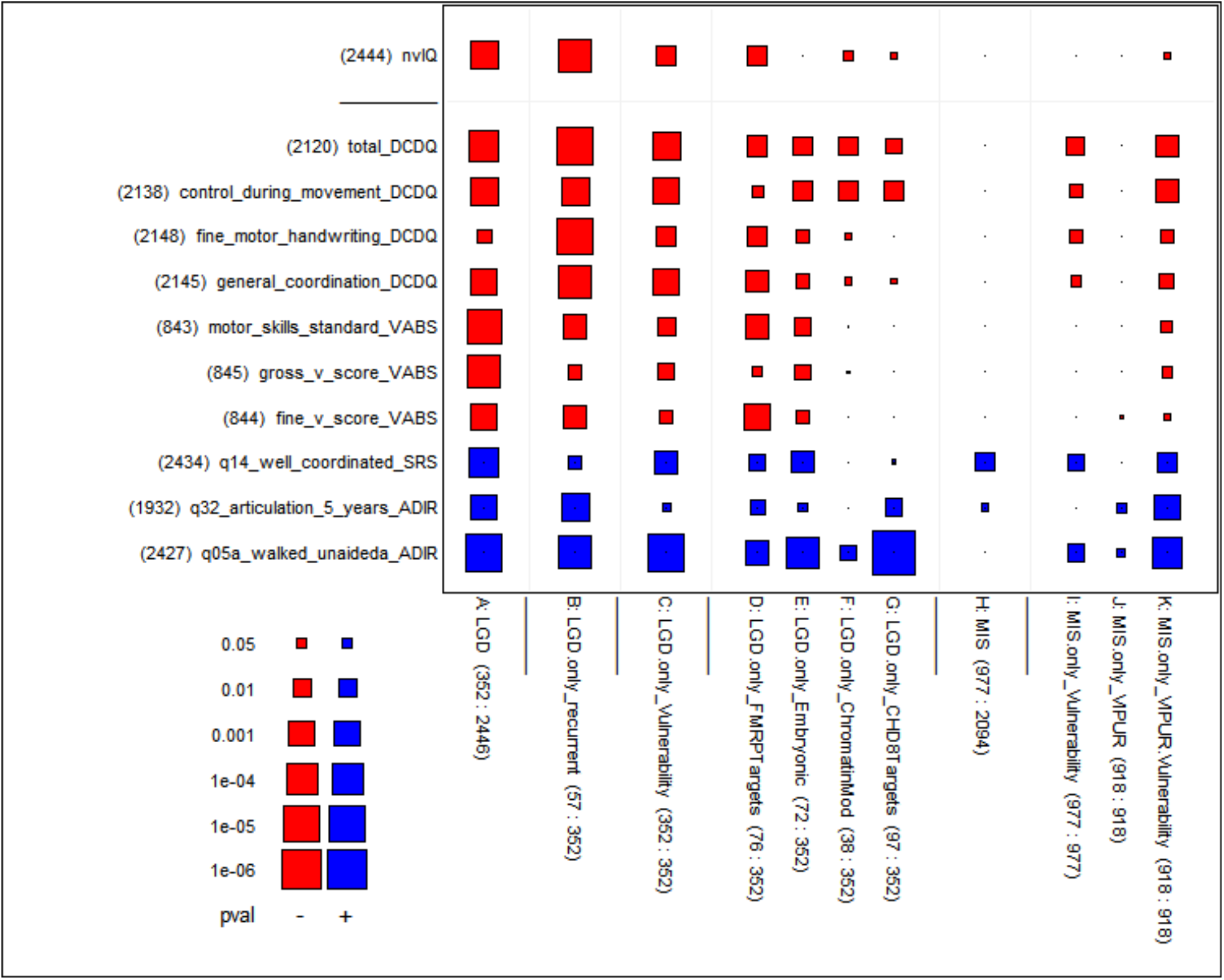
Significance of correlations between measures of genetic damage and measures of motor skills and nvIQ of affected children. We used eleven measures of genetic damage shown as columns in the figure and eleven phenotypic measures (one for nvIQ and ten for MS extracted from four different phenotypic instruments) shown as rows. The genetic damage and phenotypic measures were defined on different subsets of the affected children in the SSC collection. To reflect this fact, we pasted counts to the labels as follows: “(100)” would mean the measure is defined on a subset of size 100, and “(20:100)” indicates in addition that of the 100 values the number of non-zeroes is 20; this second version is relevant for genetic variables that are counts of mutations in a child. We computed the correlation between each of the 11 genetic damage measures and the 11 phenotypic measures using only the children for which both the genetic damage and phenotypic measures were defined, and we tested if the correlation was significantly different from 0. The resulting 11 by 11 table of p-values is rendered graphically with rectangles whose size represents inversely the p-value from the statistical test (large rectangle ≈ small p-value) and whose color represents the sign of the correlation (blue ≈ positive, red ≈ negative). The meaning of both the sizes and colors can be gleaned from the figure’s key in the bottom left. A small dot is used when the correlation is strongly insignificant (p-value ≥ 0.10). — In a similar fashion, the related Figure 1’ in the supplement shows the underlying computed correlations. For more background about this type of display, see the section Materials and Methods. **Measures of genetic damage**: The primary measure of genetic damage (column A) is defined as the number of de novo LGDs (0, 1, 2 or 3) identified in an affected child. The variables in columns B-G differentiate LGDs according to indications that they may be damaging; these variables are defined only for children who have at least one LGD. Column B is the number (0 or 1) of de novo LGDs in a child affected by genes with more than one de novo LGD in the SSC (“recurrent genes”). Column C is the sum of the vulnerability scores of the genes affected by de novo LGDs in the child. Columns D-G are defined as the number (0 or 1) of de novo LGDs that fall in four gene functional classes that have previously been implicated in autism’s etiology: FMRP target genes, embryonic genes, genes encoding chromatin modifiers, and CHD8 target genes. The remaining columns concern de novo missense mutations: Column H is the number of de novo missense mutations (0 up to 5) in an affected child, applied only to children without de novo LGD mutations, to prevent confounding with overpowering LGD effects. Column I, analogous to column C, is the sum of vulnerability scores of genes affected by missense mutations in a child. Column J is the sum of VIPUR scores of missense mutations in a child. Column K is the product of Vulnerability and VIPUR scores, exhibiting p-values that neither score could achieve alone. **Phenotypic measures**: The top row represents non-verbal IQ (nvIQ). The other rows represent ten different measures of motor skills available in the phenotypic database of the SSC. The labels are suffixed by abbreviations of the originating instruments: DCDQ, VABS, SRS and ADI-R. See Methods and Materials for details.

Because earlier literature has pointed to nvIQ as correlated with genetic lesions, we include this variable for comparison with the MS variables. Note that significant correlation with nvIQ is also seen (top left in Figure 1, p=4E-4).

### Correlating loss with LGD targets

Not all dn LGD mutations necessarily disrupt critical gene functions. In fact, we estimate that in children on the spectrum, a little less than half of dn LGDs contribute to autism-risk (Iossifov et al. 2014). First, not all LGD mutations are disruptive (recall “L” in “LGD”); and second, not all gene targets are critical. Fortunately, the importance of a gene target can be weighted by evidence. Indeed, there are multiple ways to weight target importance: whether the gene is a recurrent target; whether the gene is under strong negative selective pressure; and according to the functional properties of its encoded product.

We call a gene a ‘recurrent target’ if a dn LGD hits that gene in more than one affected child in the SSC. Such LGDs are called recurrent even though the precise dn variant itself is almost never seen twice. From previous work (Iossifov et al. 2014) recurrent targets are estimated 90% likely to be autism-risk genes. To determine if measures of MS are correlated with the presence of recurrent LGD targets, beyond their correlation with dn LGDs in general, we restricted our study to only those children already affected with a dn LGD. Among these, we then counted the number of recurrent dn LGDs in each child, typically 0 or 1. Although there are only 57 recurrent LGDs out of a total of 352 LGDs, their correlations with DCDQ in this subset of the children has very strong additional significance (Figure 1, column B, “total DCDQ”, p=5E-8). Increase in significance is found as well for nvIQ (p=8E-7), consistent with what we have previously reported (Iossifov et al. 2014).

Another way to distinguish the severity of an LGD is by characterizing the ‘genetic load’ of its target in the human gene pool, a reflection in part of the action of purifying selection. In Iossifov et al. (2015), we measured the frequency that an LGD variant in a given gene is observed in a large unaffected population. We ranked genes by their frequency of carrying an LGD in that population, normalized by the length of that gene. Those genes with a low burden were considered by us to be highly ‘vulnerable’. The data on genetic load is far from complete, because the sequence databases are insufficient to characterize most genes for their vulnerability, especially the ones encoding smaller products. Nevertheless, in a previous study we still found that affected children in the SSC with dn LGDs in highly vulnerable genes had significantly lower IQ than in affected children with LGDs in less vulnerable genes (Iossifov et al. 2015). By comparing the number of LGDs by vulnerability score in affected versus unaffected individuals from the SSC, we can demonstrate that the ability of the vulnerability score to discriminate these two groups is concentrated in higher scores (see S1 in the supplement). In this report, therefore, we transform the gene vulnerability rank by taking the negative logarithm of the normalized rank of gene vulnerability (see Materials and Methods), yielding a ‘gene vulnerability score’ that spreads out more informative scores and compresses less informative scores. Using the sum of the scores of the dn LGD target genes within a child, and restricting to children with dn LGDs, we find strikingly significant correlation with motor skills (Figure 1, column C).

The severity of a mutation might also depend on the functional class of its target gene. We consider here dn LGD mutations in three sets of genes, enriched as targets of disruptive mutation in children with ASD and earlier examined by us (Iossifov et al. 2014): FMRP target genes, whose transcripts interact with the fragile X protein (Figure 1, column D) (Darnell et al. 2011, Iossifov et al. 2012, 2014), embryonic genes, which are genes expressed in the brain of the fetus but strongly downregulated upon birth (Figure 1, column E) (Kang et al. 2011, Voineagu et al. 2011, Iossifov et al. 2014), and chromatin modulating genes (Figure 1, column F) (McCarthy et al. 2014, Bernier et al. 2014). We add a fourth set, the genes regulated by CHD8, the most frequent target gene for dn LGD mutation in ASD (Figure 1, column G) (Cotney et al. 2015). For this analysis, we again consider only those autistic individuals that have dn LGD mutation, so as not to confound the analysis with correlation due to the LGD itself. As reported before, dn LGD mutation in the FMRP target genes are associated with lower IQ (Iossifov, 2014). So, too, we find that they are significantly associated with reduced motor skills. The dn events in the other functional categories are also associated with reduced MS but with somewhat less significance. Across all the data, a particularly significant correlation is seen between presence of a dn LGD in a CHD8 target and age upon first walking. This is striking especially as the other correlations of this mutation class are weak. A related discovery was made by Bernier et al. (2014) who found that dn mutations on the CHD8 gene itself are associated with “a subtype of autism early in development.”

### Analysis of dn missense mutations

There are many more dn missense mutations than dn LGD mutations. Based on what we call ‘ascertainment bias’, we judge that only about 10% of these contribute to simplex autism, in contrast to about 50% for dn LGD (Iossifov et al. 2014, De Rubeis et al. 2014). Not surprisingly, even excluding children with dn LGDs, and counting each child for number of dn missense mutations, we see no significant correlation with nvIQ, and only significant correlation with “well-coordinated” among the MS variables (column H). Scoring for recurrent dn missense targets results in no significant correlations (not shown). However, evidence of their contribution does emerge if the dn missense mutations are weighted by gene vulnerability scores of their targets (Figure1, column I) (Iossifov et at. 2015). With that measure, we observe significant correlation to most MS variables. We note that association of these weighted mutations is not observed with nvIQ, suggesting milder effects of dn missense mutations by affecting mostly MS and leaving cognition mostly intact.

In an attempt to further strengthen the discrimination among dn missense mutations, using a method called VIPUR, we predicted the likelihood that they disrupt protein structure, and thereby infer the likelihood that they have a deleterious effect on function (Baugh et al. 2016). The two VIPUR-related variables shown in the figures are restricted to a subset of dn missense mutations for which VIPUR scores are available, aggregated by summation for each affected child without dn LGDs. The raw VIPUR score alone shows no significant correlation with MS variables (SRS item 14) and none with IQ (Figure 1, column J). However, a new score obtained by multiplication of VIPUR and the vulnerability score (see Methods) leads to more significant associations than either alone (Figure 1, column K compared to columns I or J). Thus, VIPUR in combination with the gene vulnerability score helps to assess mutational damage.

### Association between MS and IQ

It is clear from the SSC phenotype data that IQ and MS are themselves correlated (corr=0.36, N=2365). One should therefore ask if significant correlations with dn mutations survive when IQ and MS are adjusted for each other (see Materials and Methods). We display the results in Figure 2, with the same column structure as in Figure 1, where MS variables are adjusted by nvIQ, sex and age, and nvIQ (for comparison) is adjusted by total_DCDQ, sex and age. Correlations generally survive adjustment, although with somewhat attenuated significances. Attenuation after mutual adjustment is to be expected due to the partial positive correlation between IQ and MS, but the main conclusion is that the association of MS with mutation variables cannot be reduced to IQ, or vice versa. (For a corresponding display of correlations as opposed to p-values, see Figure 2’ in the supplement.)

**Figure 2.**
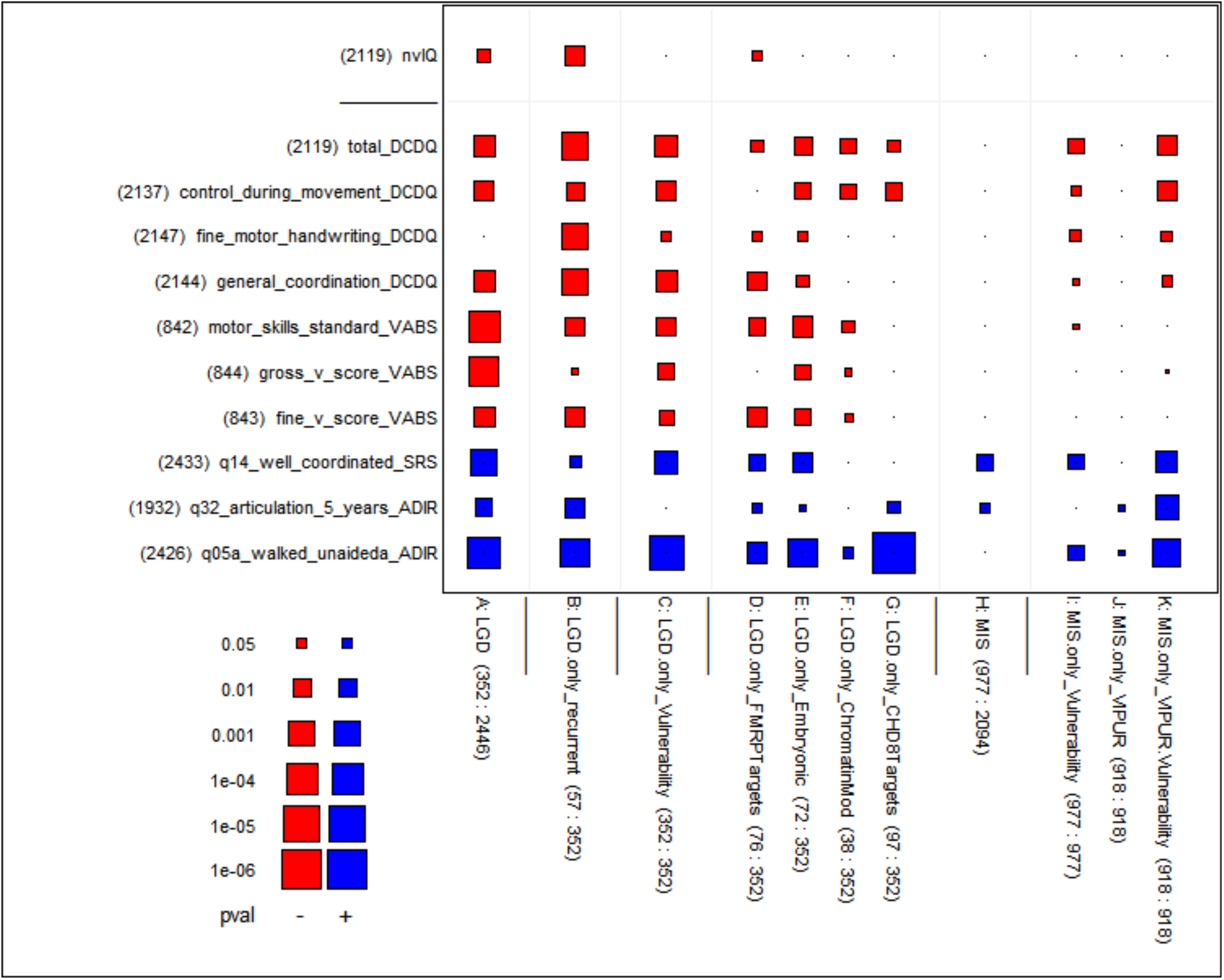
Significance of the correlation between measures of genetic damage and adjusted measures of motor skills and IQ of affected individuals. This figure is similar to Figure 1 (see its legend for details), but the phenotypic measures have been adjusted as follows: the ten motor skills measures (suffixed DCDQ, VABS, SRS, ADIR) are **adjusted for nvIQ, sex and age**; nvIQ (top row) is **adjusted for total_DCDQ, sex and age**. The main result is that the significance of the correlations largely survives these adjustments. This is evidence that the association of motor skills with the genetic variables cannot be reduced to the correlation between motor skills and nvIQ. (See Figure 2’ in the supplement for a similar graph showing the underlying adjusted correlations.)

The association between nvIQ and MS deserves closer examination because it is insufficiently characterized by a plain correlation coefficient. Indeed, the association cannot be simply described as a tendency to pair high with high and low with low values between nvIQ and MS. Rather, as can be seen in Figure 3, the combination of low nvIQ with high (normal) MS is rare, but the combination of high (normal) nvIQ with low MS is not. This can be put differently as follows:

- MS differentiate within the normal nvIQ range, while
- nvIQ differentiates within the low MS range.

**Figure 3.**
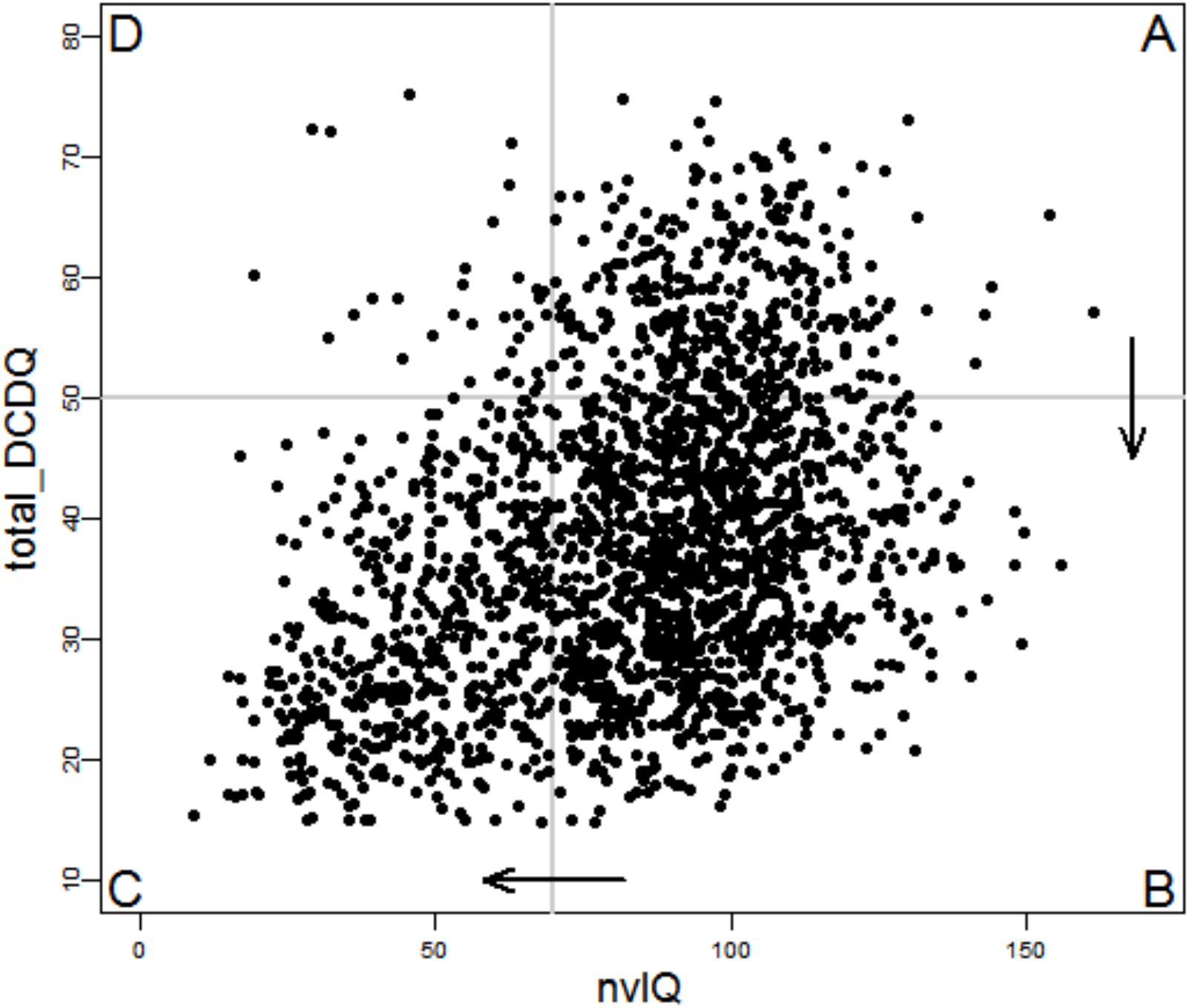
Relationship between IQ and motor skills. The scatterplot shows nvIQ and total_DCDQ for N = 2,119 affected children with available exome data. The gray vertical lines show the cut-offs used to dichotomized the two measures (see the text for justification of the particular cut-offs). The four quadrants of the graph are labeled clockwise with letters A through D. Quadrant D is significantly underpopulated compared to what is expected under an assumption that the two measures are independent. Ignoring quadrant D, the arrows demonstrate increasing phenotypic severity (as a function of both nvIQ and total_DCDQ) between adjacent quadrants, with A < B < C.

The same fact can be rendered differently by dichotomizing both nvIQ and MS: we define

- nvIQ < 70 as low IQ;
- total_DCDQ < 50 as low MS.

The value 70 for nvIQ is a conventional threshold for intellectual disability (ID), defined as two standard deviations below the mean in general populations. For total_DCDQ we use the value 50 as a loose lower bound on the “normal” range. (Following Wilson et al. (2009), the recommended age-dependent thresholds for the normal range are total_DCDQ ≥ 47 (age < 8yrs), total_DCDQ ≥ 56 (8yrs ≤ age < 10yrs), and total_DCDQ ≥ 58 (10yrs ≥ age), respectively. By this recommendation, 83% of affected SSC children have deficient MS (consistent with the 80–90% range of Hilton et al. 2012, as well as Green et al. 2009), whereas just 25.5% have diminished nvIQ (ID). Only 15.7% of affected children are in the normal range for both MS and nvIQ.)

Figure 3 gives a graphical rendition of dichotomization, resulting in four sectors denoted **A, B, C** and **D**, shown with a cross-hair at coordinates (70,50). We can order the three enriched sectors according to phenotypic severity, and label them as follows:

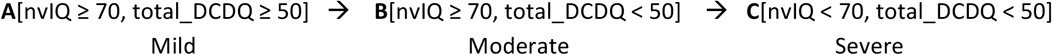

The remaining sector **D**[nvIQ < 70, total_DCDQ ≥ 50] is depleted by a factor of nearly 5 when comparing column ratios (odds ratio=4.95; see Table 2 in the supplement). In terms of severity, the phenotypic ordering of the three major sectors may be symbolically written as **A < B < C**.

Likewise, there exist significant differences between the mean vulnerability scores of LGD and missense mutation gene targets in affected individuals in Sectors **A, B** and **C**. That is, the mean vulnerability score is significantly greater

- in Sector **B** compared to Sector **A** (p=0.02 for LGDs alone and p=0.005 for LGDs and missense together);
- in Sector **C** compared to Sector **B** (p=0.02 for LGDs alone and p=0.003 for LGDs and missense together).

Therefore, in terms of ‘mutational severity’, which we use as shorthand to characterize dn mutations by the vulnerability score of the genes into which they fall, the ordering of sectors may also be written as **A < B < C**.

### Implications of the Associations between MS, IQ and Mutational Severity

We hypothesize that the increase in impairment from Sector **A** to **B** and from **B** to **C** stems from a corresponding increase in mutational severity in the target genes, and suggest the following:

- Mild mutational severity is unlikely to affect MS or nvIQ,
- moderate mutational severity is more likely to affect MS and less so nvIQ (Sector **B**), and
- severe mutational severity is likely to affect both MS and nvIQ (Sector **C**).

In other words, when the LGD or missense target has a high vulnerability score, then both diminished MS and nvIQ are more likely, and when the target has a somewhat lower vulnerability score, then the effect is more biased towards diminished MS than diminished nvIQ.

We see further evidence for the hypothesis that moderate mutational severity affects primarily MS by comparing the top two rows of Figure 1: There are no genetic variables in our collection that are more significantly associated with nvIQ than with total_DCDQ. In addition, functional classes of dn LGD mutations (other than FMRP) and the scored dn missense mutations have more significant associations to total_DCDQ than nvIQ.

### Absence of Association with Core ASD Variables

We turn to a set of core ASD variables that characterize the conventional autism phenotype consisting of deficits in social interaction as well as restricted and repetitive behaviors. In the SSC, principal investigators had previously selected “Core Descriptive Variables” from several primary instruments. From these variables we sub-selected those originating from the following instruments: Autism Diagnostic Interview (ADI-R), Autism Diagnostic Observation Schedule (ADOS), Repetitive Behavior Scale (RBS), Aberrant Behavior Checklist (ABC) and Social Responsiveness Scale (SRS, t-scores), for a total of 12 variables.

We first observe that these core ASD variables strongly correlate with both nvIQ and MS. Indeed, according to the last two columns of Figure 4, most p-values are beyond conventional levels of statistical significance. (According to Figure 4’ in the supplement, some of the correlations approach 0.5 in magnitude.) This observation suggests that core ASD variables might also be correlated with de novo mutational severity. However, transitivity of correlation is not guaranteed from statistical considerations, and so we were eager to learn directly if core ASD variables are also correlated with mutational severity. Surprisingly and importantly, we were unable to find consistent associations with the magnitude of significance seen for MS and nvIQ. As can be seen from Figure 4, strongly significant associations with genetic variables are largely absent, and some that approach the level 0.05 have the wrong sign. One exception that stands out is the variable “SRS Parent t-Score.” However, its significant correlations with genetic variables appears mediated through nvIQ, as they disappear if it is adjusted for nvIQ.

**Figure 4.**
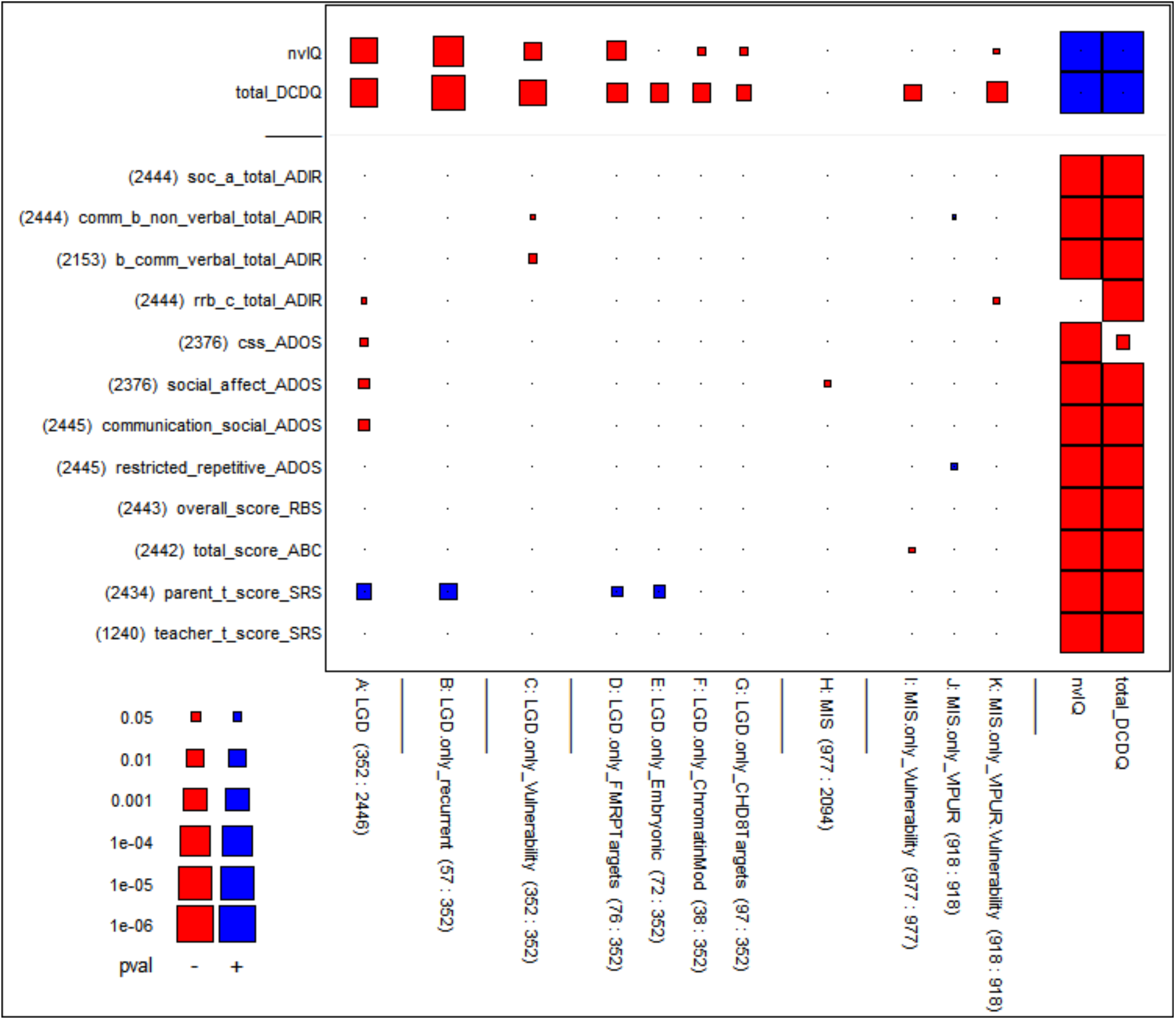
Absence of association between measures of genetic damage and measures of core ASD phenotype. In this figure, all but the top two rows represent core ASD measures drawn from the ADI-R, ADOS, RBS, ABC and SRS instruments (see Materials and Methods for explanations). The conclusion is that these measures largely lack significant correlations with measures of genetic damage from de novo mutations. The few significant correlations for parent_t_score_SRS (second row from bottom) disappear after adjustment for nvIQ (not shown). Some p-values that approach 0.05 correspond to correlations that have the wrong sign (shown red) as the core ASD variables measure behavioral deficiency, hence should be positively correlated with mutational severity. — The figure also shows nvIQ and total_DCDQ in the two top rows to provide a comparison what significant correlations would look like. The two rightmost columns show nvIQ and total_DCDQ as well to give evidence of their strongly significant correlations with the core ASD measures.

## Discussion

This study is part of our continuing attempt to link damaging de novo mutations to broad neuropsychiatric effects in children on the autism spectrum. We have done this both to define the substructure of the syndrome and to evaluate which events are likely to contribute to the disorder. Earlier studies had established a link between damaging mutation and diminished nvIQ. The present study establishes a more significant association between damaging mutation and impaired motor skills. Our method is to correlate measures of broad neurological function (nvIQ, MS) and core ASD phenotypes with de novo genetic damage. For both we rely on the SSC, which provides an abundance of phenotypic measures as well as extensive mutation data. From the latter we use information about the type of mutation (LGD or missense) and its genetic target (recurrence, vulnerability, and functional class).

Unlike intellectual ability, motor skills do not have a single standard of measurement. We therefore used all the available motor skill measures from the SSC, often broken into subscales and originating in multiple distinct instruments. Whether measuring fine or gross motor skills, or motor-related developmental milestones, with very few exceptions we see a remarkably consistent pattern of significant correlation with genetic variables. It is this consistency that gives us confidence in our conclusions.

Our present study on motor skills recapitulates conclusions from our previous study of nvIQ by extending them to motor skills (MS). First, mutational severity matters. Autistic children with LGD mutations are more likely to have impaired MS than children without them. Second, genetic targets matter. Autistic children with dn mutations in recurrent gene targets are more likely to have impaired MS. Similarly, children have more severely attenuated MS if they suffer LGD mutations in genes that are ‘vulnerable’ (i.e., have reduced deleterious genetic load in the human population) or that share certain functional properties.

This study goes beyond our previous study of damaging mutations and nvIQ. We previously had not observed correlations between nvIQ and missense mutations. The association of impaired motor skills with dn missense mutation has marginal significance at best. However, when the missense targets are weighted by their vulnerability score, the association with MS becomes far more significant. When the missense is further weighted by VIPUR, one of several available methods to judge the severity of a missense mutation, the association becomes more significant still. Importantly, while we see significant correlation between missense mutation targets weighted in this fashion and MS, we do not see it with nvIQ. As missense will be less damaging in general than the premature termination caused by LGDs, this result suggests that loss of MS is a more sensitive indicator of genetic damage than is loss of nvIQ.

nvIQ and MS as measured by the DCDQ are correlated, but far from redundant: their relation is not simply linear; they differ in gender bias (Table 1 in the supplement); they have different patterns of correlations with genetic variables; and importantly, the correlations of MS and nvIQ with genetic damage each survive when adjusting one for the other. The correlations of genetic variables with MS are more consistent than with nvIQ. Furthermore nvIQ differentiates the low functioning range of MS, while the MS differentiates the normal IQ range: moderate genetic damage tends to affect MS more and nvIQ less, while severe genetic damage tends to affect both MS and nvIQ. The most extreme example of this is the signal from children with dn LGDs in the gene targets of the CHD8 chromatin modifier. These children show very strong impairment in age of first walking, but much less significant correlation with nvIQ.

The links between damaging mutations described here should reinforce the need to routinely include an age appropriate evaluation of motor skills in the assessments of ASD. They are simple to measure. Even a single questionnaire item may give some indication of motor skill deficiency, as illustrated by SRS item 14 that refers to general motor control and coordination, and ADI-R item 32 that refers to speech articulation. Even the DCDQ instrument is relatively simple, based on just 15 items. Specific motor skills are used to define common developmental milestones for infants, one of which makes a powerful appearance in our battery of motor skill variables, the age of first walking unaided (see bottom row of Figures 1 and 2). Other related phenotypes should be examined for links to genetic damage; a promising example is dyspraxia (deficient production of gestures) studied by MacNeil and Mostofsky (2012) and Dziuk et al. (2007). In general, there is a need for more fundamental and objective tests of neurological function, such as sensory evoked responses, and tests of motor control, such as electromyography, in the routine evaluation of children with neurodevelopmental disorders.

Both nvIQ and MS significantly correlate with the core phenotypes used to make the ASD diagnosis, including those measuring social communication skills (Figure 4 and Figure 4’ in the supplement). It is natural to view nvIQ as entangled with these measures, but the entanglement with MS is less obvious. It is tempting to think of loss of MS as a consequence of broad neurological impairments, dismissing the loss as incidental to their impact on the core ASD variables. We believe this conclusion to be premature. For example, the brain region most closely associated with gross and fine motor control is the cerebellum, and it is now appreciated that the cerebellum is directly involved in cortical development and that cerebellar lesions during the third trimester and neonatal period are associated with the development of affective disorders (Wagner et al. 2017, Wang et al. 2014, Schmahmann 2013, Limperopouls et al. 2014, Bolduc et al. 2012, Messerschmidt et al. 2008).

Moreover, delay in developing age-appropriate body language may lead to further social isolation. Therefore, the connection between MS and ASD phenotypes may be direct. In any case, MS might be more readily and objectively monitored than other clinically emphasized cognitive functions, and so might be an important endpoint while investigating genetic lesions in model organisms or while screening humans for response to experimental therapies.

Finally, we examined the correlation between damaging mutations and the core ASD variables, including social deficits and restricted and repetitive behaviors. Although damaging mutations correlate with both nvIQ and MS in those with ASD, and nvIQ and MS correlate very significantly with core phenotypes, it does not follow that damaging mutations will correlate with core ASD. The correlations of mutations with nvIQ and MS, while significant, are weak in absolute terms, and transitivity in correlation is not forced. In fact, we find that the correlation of damaging mutation with core ASD is inconsistent and weak at best. The little correlation observed vanishes when we adjust for nvIQ.

The lack of correlation, nevertheless, merits speculation. Our first thought was that this absence of significant association could be explained by the use of some core ASD variables in the ascertainment of children with ASD in the SSC, causing truncation of the variable ranges and resulting in less significant associations. A closer examination showed, however, that this explanation is most likely wrong (see S2 in the supplement). We can consider another explanation, one we cannot rule out. Restating our result, we can say that whereas damaging de novo mutations are strongly associated with the diagnosis of ASD (see for example Iossifov et al. (2014), Figure 1), they are not consistently correlated with core ASD variables among affected children. Thus, if social deficiencies and restricted and repetitive behaviors also have contributing genetic causes, perhaps they arise from shared ancestral variants transmitted from parents. These variants may not be under negative selection, and hence persistent and not very rare in the population. Rather than causing deleterious phenotypes in their own right, these variant alleles may be responsible for the diversity of human social, communication and cognitive traits that characterize the human species.

### Materials and Methods

The present study is based on the Simons Simplex Collection (SSC, Fischbach and Lord 2010), which has data for 2760 families that have a single child affected by ASD. Of these families, 2280 have an unaffected child as well, but for the most part, we are only concerned with the affected children. Among them, 2446 have exome sequencing data available that resulted in the identification of 3403 de novo mutations of all types (Iossifov et al. 2014). (The 1836 unaffected children with exome data have 2288 identified de novo mutations among them.) Unlike case-control approaches that compare affected and unaffected children, ours is a study of association between phenotype and genotype variables among affected children only. The premise, which could have been wrong, is that the affected children in the SSC have sufficient phenotypic variation to allow the discovery of statistically significant correlations between high and low levels of a scored behavioral phenotypes (such as nvIQ, MS and core ASD variables) on the one hand, and genotypic events (such as different classes of de novo mutation and the characteristics of their target genes) on the other hand. As shown above, convincing correlations exist for nvIQ and MS variables, but not for most core ASD phenotypes.

The following is the list of MS variables shown by their names in the SSC tables which also convey the intended meaning of the scales:

- **DCDQ** (Developmental Coordination Disorder Questionnaire):

- control_during_movement
- fine_motor_handwriting
- general_coordination o total The last is the summary scale; the preceding three are subscales formed from a pool of 15 items scored on a 5-point Likert scale. Orientation: high values for high achievement.
- **VABS-II** (Vineland Adaptive Behavior Scales – II):

- fine_v_score
- gross_v_score
- motor_skills_standard Again, the last is the summary scale; the preceding two are subscales. These variables exist only for children up to age 7.5 years (cases with higher age are outliers that were removed). Orientation: high values for high achievement.
- **SRS** (Social Responsiveness Scale):

- q14_well_coordinated This is a single item out of 65 SRS items that measures coordination problems on a scale from 0 to 3. Contrary to the meaning suggested by the name of the item, this is a severity measure with meanings 0 = “no coordination problems”, 3 = “severe coordination problems”.
- **ADI-R** (Autism Diagnostic Interview – Revised):

- q05a_walked_unaideda: *Age* of walking unaided, in months (ranging from 7 to 72), but transformed with a double logarithm due to an extremely right skewed distribution: log(log(…)).
- q32_articulation_5_years: Problems with motor control of speech at age 5, on a scale from 0 to 3 Orientation: high values for higher level of problems. Note: Consistency of correlation across diverse measures of MS from multiple instruments strengthens our confidence that the conclusions are not measurement artifacts.

**Cognitive functioning** is measured by non-verbal IQ (nvIQ), in agreement with past literature. Verbal and full-scale IQ are not used.

**Core-ASD variables** were selected from the SSC table “Core Descriptive Variables” (CDV), which contains a set of demographics, measures and diagnoses previously deemed clinically relevant. We subselected 12 variables from 5 instruments as follows:

- **ADI-R** (Autism Diagnostic Interview – Revised): adi_r_soc_a_total, adi_r_comm_b_non_verbal_total, adi_r_b_comm_verbal_total, adi_r_rrb_c_total
- **ADOS** (Autism Diagnostic Observation Schedule): ados_css, ados_social_affect, ados_communication_social, ados_restricted_repetitive
- **RBS-R** (Repetitive Behavior Scale – Revised): rbs_r_overall_score
- **ABC** (Aberrant Behavior Checklist): abc_total_score
- **SRS** (Social Responsiveness Scale): srs_parent_t_score, srs_teacher_t_score

These variables were pre-selected on substantive grounds without datamining. However, the interested reader may indulge in “datamining” by perusing the numerous figures in the supplement (S4-S8) which shows associations for the complete instruments, both summary measures and underlying items. These figures are provided as exploratory displays and as confirmations and qualifications of the finding that the core ASD phenotype has at most a tenuous link to genetics as reflected by de novo mutations. The instruments we show in the supplement are as follows: **DCDQ, VABS-II, CDV** (SSC Core Descriptive Variables), **CUV** (SSC Commonly Used Variables), **CBCL-2–5** and **CBCL-6–18** (Childhood Behavior Checklist, ages 2–5 and 6–18), **SRS, ABC, RBS-R, ADI-R, ADOS-1, ADOS-2, ADOS-3** (three modules of **ADOS**), **SCQ-LIFE** (Social Communication Questionnaire – Life), as well as a table of demographic variables.

**Genetic variables** for exome-sequenced affected children were obtained from published sources as follows:

- The list of de novo mutations is from supplementary Table 2 in Iossifov et al. (2014). It characterizes each mutation by the “location” on the genome, the “effectGene”, and the “effectType” (among other things). Among effectTypes we used the following: “synonymous”, “missense”, as well as six types that jointly make up the LGD classification: “splice-site”, “nonsense”, “noStart”, “noEnd”, “frame-shift”, “no-frame-shift-newStop”.
- The functional classification of genes is from supplementary Table 7 in Iossifov et al. (2014). LGD mutations where subdivided according to whether the “effectGene” is classified as “FMRPTargets”, “Embryonic” or a “ChromatinModifiers” (remaining classifications were not used, some because of low counts, others because of *a priori* unlikely effects).
- One more classification was used to subdivide LGD mutations according to their “effectGene”: “CHD8Modifiers”, comprising a list of genes published by Cotney et al. (2015), supplement 1.
- Gene vulnerability ranks are from Iossifov et al. (2015). Instead of dichotomizing the scores on a threshold and comparing the resulting groups of high and low gene vulnerability, we instead transform the ranks by normalizing them to values between 0 and 1, and then applying a negative logarithm. The resulting “gene vulnerability scores” have a roughly exponential distribution, the purpose being to spread out the most vulnerable genes to the unlimited positive range and shrinking non-vulnerable genes to the near-zero range. This processing gives highly vulnerable genes an opportunity to differentiate themselves with high values while genes with little vulnerability are made nearly indistinguishable by piling up their values near zero. This type of processing injects quantitative differentiation where it is needed and obviates the search for meaningful thresholds.
- VIPUR scores were obtained from the authors of Baugh et al. (2016). We used the “highest score” version of VIPUR. Similar to gene vulnerability scores, we did not use the raw VIPUR scores but a transformation thereof, again obtained by normalizing their ranks to [0,1] and applying a negative logarithm, resulting in a roughly exponential distribution that spreads out the high scores and shrinks the low scores.

**Descriptive tables and plots** for nvIQ, MS variables and genetic variables are shown in Section S9 of the supplement. Similar information for the many instruments shown in Sections S3-S8 are not provided due to shear volume.

**Statistical measurement of association** between two variables was done by forming Pearson correlation coefficients. These were used for uniformity even in non-standard cases, in particular when one or both variables were binary groupings coded as 0–1 dummy variables: If one variable is quantitative and the other binary, the correlation coefficient is algebraically equivalent to the t-statistic for testing the difference of means; if both variables are binary, the correlation coefficient is algebraically equivalent to the test statistic of Fisher’s exact test of independence. See Buja et al. 2014 for a more detailed discussion.

**Presentation of correlation tables** is in graphical form as “blockplots” (Buja et al. 2014). The reason is that large tables of numeric values are difficult to parse visually. Furthermore, the multi-digit precision of numeric tables is not only useless but delusional because it suggests accuracy where none exists. Graphical presentation provides not only defensible accuracy but lends itself to visual pattern recognition, in particular patterns of consistency of correlations across rows and columns. Actually, more important than correlations are their statistical significances in terms of p-values. The most important figures are indeed blockplots of p-values, rendered such that large blocks indicate strong statistical significance. The visual estimation of the order of magnitude of p-values (as well as correlations) is helped by a key in the bottom left of the figures. As for color coding, we use blue to indicate a positive association and red a negative association; color coding is also used in blockplots of p-values even though these only reflect statistical significance without orientation. Finally, we note that the blockplots are superior to heat maps because the sizes of rectangles provide much more precise and more impactful visual cues than color scales.

**Statistical multiplicity**: Presenting large numbers of correlations and their p-values might raise questions of statistical inference. A more conventional presentation would have condensed the findings into a handful of p-values, for example, by focusing on total_DCDQ alone among MS variables. The reason for choosing an expansive visual presentation of large numbers of p-values is to convey the consistent patterns of statistically significant association across groups of variables, in particular the several measures related to motor skills. Such consistency is non-trivial: While motor skill variables are correlated with p-values beyond conventional levels of statistical significance, the correlations are far from perfect, no higher than 0.5 between DCDQ and VABS-II, for example (see S9.4 in supplement). This limits their shared variation to 25%, leaving ample room for conflicting correlations with the comparatively weak signal from the mutation variables. That this is largely not happening is confirmation of the non-trivial consistency of association between motor skill and genetic variables. The many p-value displays in the supplemental sections S4-S6 are shown for two reasons: 1) to back up and qualify the notion that a vast majority of core ASD variables does not show consistent correlations with mutation variables, other than those mediated by nvIQ and MS, and 2) to allow readers to do their own exploratory hypothesis generation, using p-values heuristically rather than inferentially.

**Adjusted variables**: In Figure 2 we (1) “adjust” nvIQ for total_DCDQ, sex and age, and (2) we conversely adjust the MS variables for nvIQ, sex and age. Adjustment means in case (1) that nvIQ is subjected to a linear regression with total_DCDQ, sex and age as regressors and that nvIQ is then replaced by its residuals from this regression. These residuals are uncorrelated with total_DCDQ, sex and age. If one observes correlations of “adjusted nvIQ” with genetic variables, it reflects association that cannot be accounted for by total_DCDQ and/or sex and/or age. (Detail: In adjusting for age, we added a linear spline term with knot at 9 years of age to account for potential nonlinearity due to the transition from childhood to adolescence. The knot location 9 was chosen a priori, not by data mining.)

**Permutation Tests Related to the Ordering of the Sectors “A < B < C”**: In the comparison **B > A** we are only interested in the association between vulnerability and total_DCDQ, not nvIQ. However, there exists a weak positive association between total_DCDQ and nvIQ even in the union of sectors B and A. In order to filter out the confounding with nvIQ, we performed a conditional permutation test that permutes only within deciles of nvIQ. In the comparison **C > B** the roles of total_DCDQ and nvIQ are reverse and hence permutations are within deciles of total_DCDQ only.

**Questions of Confounding**: Observational studies such as the present one can result in flawed attribution of cause. While we formulate all results in terms of association rather than causation, the implied understanding is that the genetic variables describe aspects of causal mechanisms for the phenotype. In order to reduce the chance of confounding of the genetic variables with trivializing factors such as demographics, we provide in the supplemental section S7 p-value displays for demographics, and in S8 displays for genders separately and without non-Caucasian ethnicities that could potentially confound with genetic variables. From the demographics in S7 we learn that, for example, IQ is much more associated with demographics than MS (as measured by total_DCDQ), thus reducing the chances of confounding for the latter. We also learn the known fact that dn missense mutations are more related to fathers’ age than mothers’. From the subset analysis in S8 we learn that the major findings persist when considering white males only. Due to the gender imbalance of autism, females constitute a much smaller subset in which statistical significances are necessarily attenuated, with notable exceptions for LGDs in the young females described by the Vineland (VABS) MS measures. Overall the conclusion is that non-Caucasian ethnicities and female sex are not confounding factors behind the associations between mutational severity and MS.

## Acknowledgements

We thank Jim Simons for prompting the research and providing valuable guidance throughout; Gerry Fischbach for a detailed review of the manuscript and suggestions for improvements; Louis Reichardt for providing additional pointers; and Richard Bonneau and Evan Baugh for explanations of the VIPUR method and sharing missense mutation predictions. We also thank all the families at the participating SSC sites, as well as the principal investigators (A. L. Beaudet, R. Bernier, J. Constantino, E. H. Cook, Jr., E. Fombonne, D. Geschwind, D. E. Grice, A. Klin, D. H. Ledbetter, C. Lord, C. L. Martin, D. M. Martin, R. Maxim, J. Miles, O. Ousley, B. Peterson, J. Piggot, C. Saulnier, M. W. State, W. Stone, J. S. Sutcliffe, C. A. Walsh, and E. Wijsman) and the coordinators and staff at the SSC sites for the recruitment and comprehensive assessment of simplex families, and the Simons Foundation Autism Research Initiative (SFARI) staff for facilitating access to the SSC. This work was supported by SFARI Grants SF235988 (to M.W.), SF362665 (to I.I.) and SF448357 (to A.B. and A.K.).

